# Natural selection and maladaptive plasticity in the red-shouldered soapberry bug

**DOI:** 10.1101/082016

**Authors:** M L Cenzer

## Abstract

Natural selection and phenotypic plasticity can both produce locally differentiated phenotypes, but novel environments or gene combinations can produce plasticity that works in opposition to adaptive change. The red-shouldered soapberry bug (*Jadera haematoloma*) was locally adapted to feed on the seeds of an introduced and a native host plant in Florida in the 1980s. By 2014, local differentiation was lost and replaced by phenotypically similar populations all adapted to the introduced host, likely as a result of gene flow. Here, I quantify the effects of these two host plants on individual performance, natural selection, and phenotypic plasticity. I find that the seed coat and seedpod of the native host have strong negative effects on juvenile survival and adult reproduction compared to the introduced host. I find support for the hypothesis that the seedpod is driving diversifying natural selection on beak length, which was previously locally adapted between hosts. I also find maladaptive plasticity induced by host plant: bugs develop beak lengths that are mismatched with the seedpod size of the host they are reared on. This plasticity may be the result of gene flow; hybrids in the 1990s showed the same pattern of maladaptive plasticity, and plasticity is stronger in the present in areas with high gene flow. Although ongoing natural selection has produced locally adapted genotypes in soapberry bugs, maladaptive plasticity has masked the phenotypic difference between populations in the field.

## Introduction

The role of plasticity in natural selection and local adaptation is controversial (West-Eberhard 2003; Crispo 2007, 2008). In some cases, plasticity may facilitate an evolutionary response, either by allowing populations to persist and maintain genetic variation in novel habitats or by bringing populations close enough to an adaptive peak for selection to act on underlying genetic variation (Price et al. 2003; Yeh and Price 2004; Lande 2009). In other cases, plasticity inhibits genetic change by masking genetic variation in phenotypes under selection (Sultan and Spencer 2002; Borenstein et al. 2006). Plasticity may also be maladaptive and move phenotypes away from local fitness maxima. Maladaptive plasticity may generate phenotypic differences in nature in the absence of divergent selection (eg, countergradient variation; Conover and Schultz 1995). In systems with divergent natural selection between habitats, maladaptive plasticity may dampen, mask, or even reverse the phenotypic signal of adaptive genetic differentiation in nature.

Maladaptive plasticity is expected to arise most often in novel habitats due to a lack of evolutionary history between environmental cues and physiological responses. Individuals may develop traits that would have been adaptive in the historical context, but are not adaptive in the new environment; novel habitats may also expose cryptic genetic variation, producing phenotypes that have never been exposed to selection in any habitat (Ghalambor et al. 2007). This type of maladaptation should be purged by selection relatively quickly. Novel genotypes, like those produced in hybrids or backcrosses between diverged populations coming into secondary contact, have also not experienced selection and should therefore be more likely to harbor maladaptive variation. Unlike in novel environments, however, it may be difficult for selection to purge maladaptive phenotypes in hybrids if opposing selection continues to act on each parental type.

Local adaptation to different host plants is common in plant-feeding insects (eg, Bush 1969; Carroll and Boyd 1992; Ferrari et al. 2008; Downey and Nice 2011), where host plant defenses are a major driver of natural selection and differentiation (Ehrlich and Raven 1964; Wheat et al. 2007; Toju 2009). Natural selection is often invoked as the source of local adaptation; however, other processes commonly produce local trait differentiation (eg, plasticity, correlated selection, genetic drift). To lend support to the hypothesis that a trait is under divergent selection by an environmental feature that differs between two habitats, we should first answer some basic questions: Does the proposed selective agent have a substantial impact on individual performance? Do some trait values have higher performance than others when challenged by the selective agent, and does the effect of the trait value on performance differ between environmental types? What aspect(s) of performance are influenced by the selective agent and are they important for survival and reproduction in nature?

The red-shouldered soapberry bug *Jadera haematoloma* (Hemiptera: Rhopalidae) is a textbook case of rapid local adaptation to a novel environment. Soapberry bugs are native to the southern peninsula and Keys of Florida, where they have evolved with a native balloon vine (*Cardiospermum corindum*). In the 1950s, the Taiwanese golden rain tree (*Koelreuteria elegans*) was widely introduced to the peninsula of Florida and colonized by soapberry bugs. The native *C. corindum* has large, inflated seedpods, hard seeds, high nitrogen content, and asynchronous fruiting throughout the year. The invasive *K. elegans* has flattened seedpods that are open for most of the development of the seed, soft seeds, high lipid content, and a single, synchronous fruiting period each autumn (Umadevi and Daniel 1991; Carroll and Boyd 1992; Carroll et al. 1998). By 1988 there was clear evidence of local host-associated adaptation in soapberry bugs. Juveniles had higher survival and shorter development time on their local host than on the alternative host (Carroll et al. 1997, 1998). Differences in dispersal morphology were found between host-associated populations: individuals from *K. elegans* had higher frequencies of non-dispersive morphs than individuals from *C. corindum*, hypothesized to be due to differences in host plant reproductive synchrony (Carroll et al. 2003). Adult feeding morphology was locally adapted: adult females adapted to *C. corindum* had long beaks, while females from populations on *K. elegans* had evolved short beaks (Carroll and Boyd 1992). It was hypothesized that long-beaked females would be more successful at accessing seeds inside of the large pods of *C. corindum,* which should increase nutrient acquisition and therefore energy available to invest in reproduction. On *K. elegans*, it was hypothesized that short beaks would be more efficient for feeding through the flat pods, which would increase the speed of nutrient acquisition and give females with short beaks a competitive advantage. Beak length was highly heritable (Dingle et al. 2009) and only weakly affected by developmental host plant in pure host races. In contrast, individuals with mixed ancestry showed a profound, potentially maladaptive influence of developmental host on beak length: female crosses and backcrosses between host races developed longer beaks when raised on the seeds of the small-podded host, *K. elegans*, than on the large-podded *C. corindum* (Carroll et al. 2001). All of these locally adapted phenotypes suggest divergent selection was acting between the two host plants. The direct effects of host plant defenses on performance and the role of the seedpod in natural selection on beak length have remained untested.

By 2014, the pattern of local adaptation between hosts had shifted dramatically. Adult morphology, survival, and development time are all converging across populations on both host plants (Cenzer 2016). The phenotypes that now dominate across the state of Florida are those adapted to the non-native host, *K. elegans*. It is likely that this phenotypic pattern is the result of gene flow increasing from populations adapted to *K. elegans* to those on *C. corindum*. The novel host has increased in frequency and the native host has become less common during the last 25 years (Carroll and Loye 2006). As gene flow increases, more hybrids should be present in nature, particularly on the native host. Dingle et al 2009 (using bugs collected from the field in 1991) showed that experimentally produced hybrids had higher plasticity than pure host races in response to developmental host. As gene flow increases hybridization in the field, therefore, developmental plasticity induced by each host plant is likely to become more common.

In this study, I addressed three primary questions. In order to understand how host plants drive natural selection, we need to know what plant traits have the greatest effects on insect fitness. Therefore, I first asked: 1) How do two physical host plant defenses, the seed coat and the seedpod, influence patterns of juvenile and adult performance on each host plant? I then addressed the key questions: 2) How does the host seedpod influence natural selection on beak length? and 3) How does host plant influence plasticity in morphology?

Based on known differences in physical defenses and current and historical patterns of local adaptation, I proposed the following hypotheses: H1) Juvenile mortality is driven by the tough seed coat, which prohibits early feeding and causes starvation. H2) Juveniles reared on the native *C. corindum* will develop shorter beaks than those raised on *K. elegans*, consistent with historical patterns of maladaptive plasticity documented in hybrids between host races; this change may be coupled with plasticity in body size and dispersal morph due to correlations in these three phenotypes. Alternatively, selection may have purged deleterious plasticity in areas where gene flow is high, in which case we should observe neutral or adaptive plasticity. H3) If maladaptive plasticity is being generated by gene flow, I expect a latitudinal gradient such that developmental plasticity is strongest in the south, where gene flow between hosts is more common, and becomes weaker moving north away from the sympatric zone. H4) The presence of the seedpod reduces adult fitness by reducing access to seeds. H5) Adults with longer beaks will access more seeds within closed pods and adults with short beaks will feed more efficiently on seeds in open pods, consistent with divergent natural selection on beak length.

To test these hypotheses, I conducted three experiments. The first tested the effect of the rearing host and the seed coat on juvenile survival, development time, and morphology (H1, H2, & H3; Fig. 1a). The second and third experiments tested the effect of the seedpod on adult performance and natural selection on beak length in the lab and field, respectively (H4 & H5; Fig. 1b,c).

**Figure 1.**
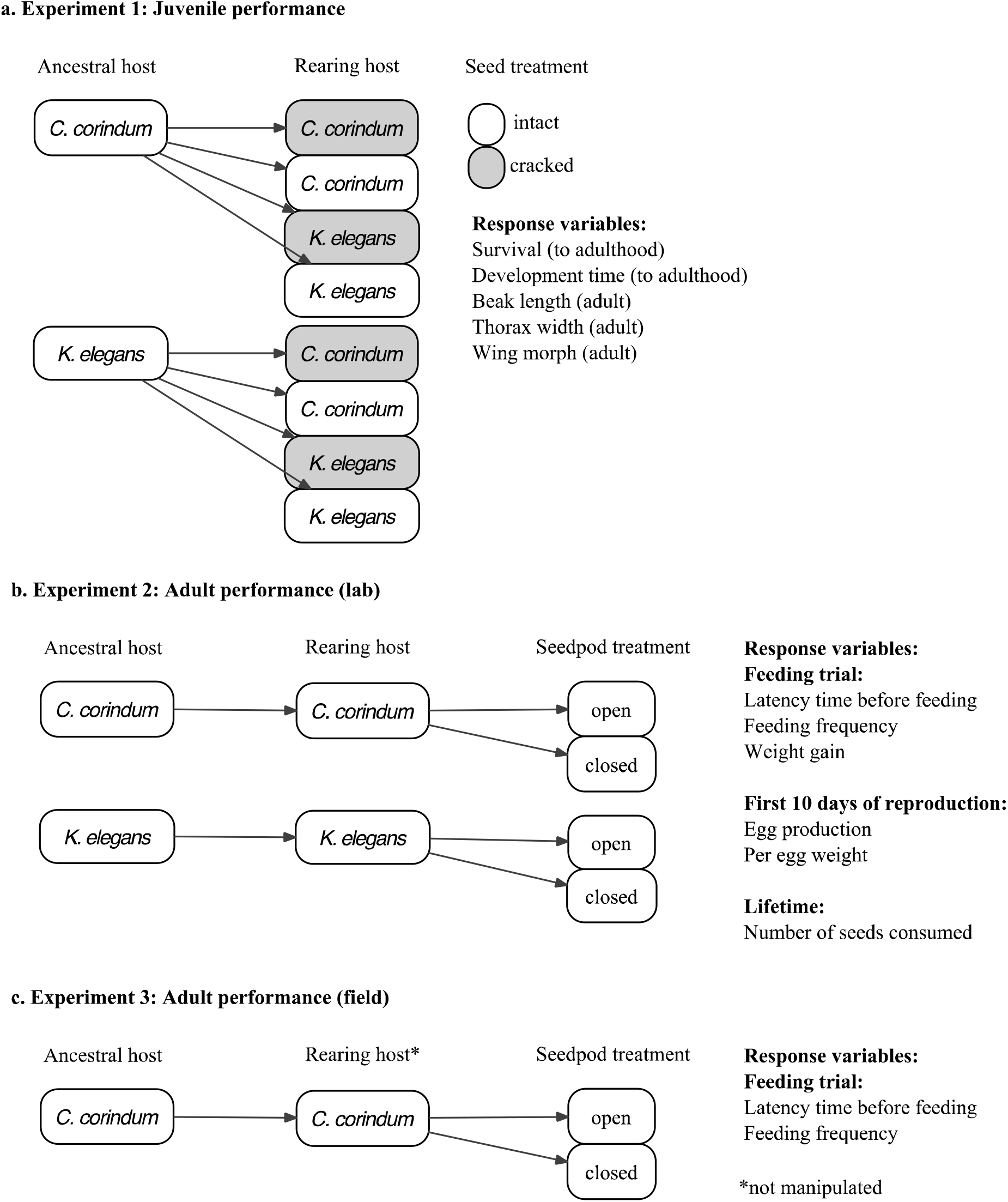
Experimental design and response variables for each experiment. a. Experiment 1 testing the effect of rearing host and the seed coat on survival, development time, and adult morphology; b. Experiment 2 testing the effect of the seedpod on adult feeding and reproduction in the lab on both hosts; c. Experiment 3 testing the effect of the seedpod on adult feeding in the field on *C. corindum*.

## Methods

### Collection

I collected *J. haematoloma* in April 2014 for the juvenile performance experiment and in December 2013 for the adult performance laboratory experiment from 8 locations in Florida (Fig. 2; coordinates in Table S1). The adult performance field experiment was conducted in April 2014 with field-collected individuals from Key Largo. Populations at the northern four sites (Gainesville, Leesburg, Lake Wales, and Ft. Myers) occur on *K. elegans* and the southern two sites (Key Largo and Plantation Key) occur on *C. corindum*. Bugs occur on both host plants in Homestead, the only known location where these two host plants occur in close proximity (<5km). For laboratory experiments, I collected host plant seeds from each site in December 2013 and April 2014 and stored them at 4°C until they were used for rearing. I discarded seeds with visible indications of previous feeding and tested all seeds for viability by placing them in water and discarding seeds that floated. I collected from 5-10 individual trees at each *K. elegans* site and 3-15 individual vines at each *C. corindum* site. For the field experiment, fully inflated green pods were collected from the field and stored in gallon Ziploc bags with a paper towel for 1-2 days at room temperature before the experiment began. Pods with signs of feeding by other arthropods (especially caterpillars of the silver-banded hairstreak, *Chlorostrymon simaethis*) were discarded. Pods were gently squeezed to locate holes where first instar caterpillars may have entered if no obvious signs of feeding (eg, frass) were located.

**Figure 2.**
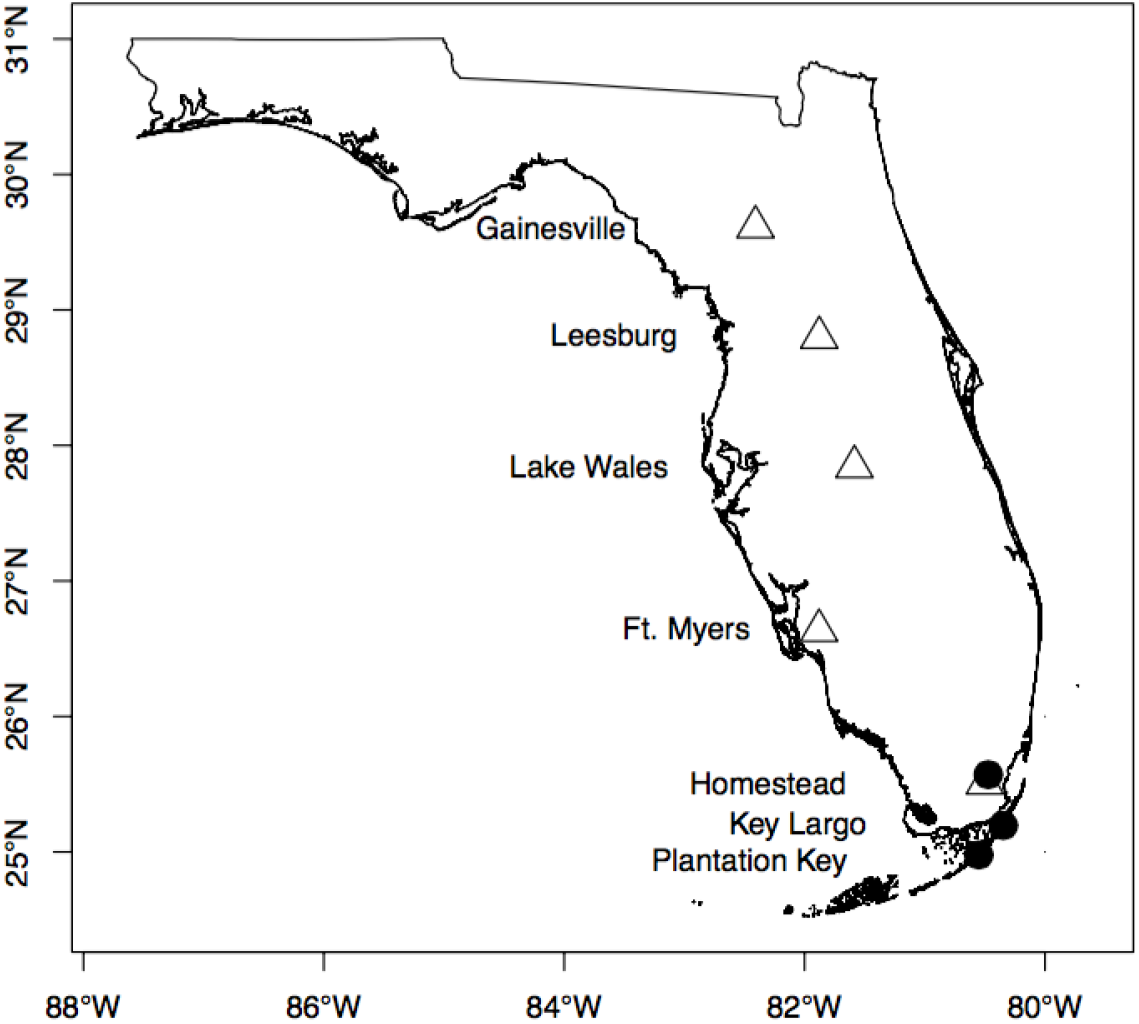
Collection sites on the introduced host, *K. elegans* (white triangles) and the native host, *C. corindum* (black circles) with location names. Sites are abbreviated in text as follows: Gainesville=GV, Leesburg=LB, Lake Wales=LW, Ft. Myers = FM, Homestead [*C. corindum*]=HS1, Homestead [*K. elegans*] = HS2, Key Largo=KL, Plantation Key=PK.

### Rearing conditions

For laboratory experiments, all rearing was carried out in Sanyo Versatile Environmental Test Chambers at 28°C during the day and 27.5°C at night, 50% relative humidity with a 14:10 Light:Dark cycle, following spring climate conditions in the field and those used in Carroll et al 1998. Adults collected from the field were housed as mating pairs in vented Petri dishes lined with filter paper and given water in a microcentrifuge tube stoppered with cotton (“water pick”) and 3 seeds of their field host plant. Eggs were collected daily until hatching. Nymphs were removed within 12 hours of hatching to reduce egg cannibalism and housed individually in mesh-lidded cups lined with filter paper with a water pick and a seed of their assigned host plant treatment. Additional seeds (a total of 2 for *K. elegans* and 3 for *C. corindum*, for a total seed mass of ~150mg) were added at 7 days of age. This seed mass has been shown in previous experiments to be more than sufficient for individuals to reach adulthood, and is considered *ad libitum* (Carroll et al. 1997). Individual containers were rotated daily within mesh boxes (each holding 36 individuals), and boxes were rotated daily within the growth chamber to reduce the effect of specific location. Water, paper and cotton were changed weekly for nymphs and adults. Nymph survival and whether or not they had reached adulthood were assessed daily.

### Assessing the effects of rearing host and seed coat on juveniles

The first experiment (Fig. 1a) tested the effects of the rearing host species and the seed coat on juvenile survival, development time, and morphology (H1, H2, & H3). For this experiment, I used the second laboratory generation (F2) of juveniles descended from the April 2014 field collection. Their parents, the first laboratory generation (F1), were reared using the rearing conditions described above and were mated with individuals from the same population but different families to avoid inbreeding. F2 individuals were reared in a split-brood cross-rearing design: full siblings from each family were randomly assigned to a rearing host (either *C. corindum* or *K. elegans*) and a seed treatment (seeds were either intact or had the seed coat cracked) such that all families were represented in each treatment combination. The seed treatment was administered by gently clamping seeds in pliers and tightening the pliers just until a crack formed in the seed coat. Upon reaching adulthood, individuals were frozen and stored at −20°C for morphological analyses. Beak length (the distance from the anterior tip of the tylus to the distal tip of the mouthparts), thorax width (at the widest part of the pronotum), and forewing length (from anterior to distal tip) were measured for all individuals to the nearest 0.02 mm using Mitutoyo digital calipers for. Each individual was identified as either macropterous (long forewings) or brachypterous (truncated forewings)(Carroll et al. 2003).

### Assessing the effects of the seedpod on adult performance

#### Lab feeding trials

The second experiment (Fig. 1b) I conducted was designed to test the effect of the balloon vine seedpod on adult performance (H4) and natural selection on beak length (H5). For this experiment, I used adult females from the first laboratory (F1) generation descended from individuals collected in the field in December 2013. Individuals were reared following conditions described above, on the same host species their parents were collected from in the field. When individuals reached adulthood, females were isolated with water and no seeds for 2-5 days before feeding trials began to allow their cuticle to harden completely. Males were not used in feeding trials, and were given 1-2 additional seeds within 24 hours after eclosion. Males were not used in the feeding trails because they are substantially smaller than females and never have beaks long enough to reach seeds within seedpods. Furthermore, local adaptation in beak length has not been observed in males, and so there is no reason to suspect differential selection is acting on male beak length (Carroll and Boyd 1992).

Seedpods were prepared by opening dried, intact seedpods of either *C. corindum* collected in the Florida Keys or *C. grandiflorum* collected in Davis, California. *C. grandiflorum* is a congener of *C. corindum* with pods of similar shape and size that is commonly fed on by *J. haematoloma* in Davis. The average pod radius used in this experiment was 7.60±0.31mm. The mean pod size is within one standard deviation of the mean beak length measured in the field in Florida in December 2013 (7.45±0.10mm), and should therefore encompass the range of pod sizes where soapberry bugs are most likely to feed in nature. The seedpods used in this experiment were smaller than historical *C. corindum* seedpod sizes, which should make the assessment of the fitness costs of the seedpod conservative, since on average more females will be capable of reaching the seeds within the closed seedpods in the experiment than in the average naturally occurring seedpod. Both dry and green seedpods occur in nature; seedpods begin drying when the seed reaches maturity and seeds may remain inside dry pods for several weeks. It would have been interesting to compare both green and dry pods in the laboratory experiment, but it was not logistically feasible; therefore, green pods were evaluated separately in the field (described below). In laboratory experiments, the original seeds in experimental pods were removed and discarded. Three seeds of the host species from which individuals were collected in the field (*C. corindum* or *K. elegans*) were glued into the natural position and orientation within each pod with Elmer’s glue and allowed to dry for 1-2 hours. For closed pod treatments, the three segments of the pod exterior were then reattached with glue. For open pod treatments, each segment of the pod exterior was lined with glue, placed in the ‘closed’ position, and then detached again. This was done to ensure that an equal quantity of glue was present on pods of both treatment types to control for possible deterrent effects of glue on feeding. All parts of the pod were retained together and included in the treatment container. Glue was allowed to dry overnight (12-18 hours) before feeding trials were conducted.

After their cuticles hardened, females were held in mesh-lidded containers (BugDorms, MegaView Science, Taiwan) with seeds and seedpods of their seedpod treatment and a water pick for feeding observations. When soapberry bugs feed, they drill a hole into the seed using the serrated ends of their beak (modified labium). Once the hole has been created, the second and third segments of the labium bend characteristically and a thread-like stylet is inserted into the seed. For this experiment, we included both drilling behavior and stylet insertion as feeding behavior. Probing the seed without beginning drilling was not considered feeding. It was not possible to see how the beak interacted with the seed inside of closed pods; therefore, feeding was defined for bugs in the closed pod treatment as having the beak fully inserted into the pod and remaining stationary. It is not known whether soapberry bugs can satiate within 8 hours, although individual feeding bouts often last several hours. The amount of time each individual spent in the feeding trial was not a significant predictor of whether or not an individual was observed feeding, suggesting satiation was not reducing feeding activity during the feeding trail.

I conducted feeding trials of 5.75-8 hour duration in the lab at ambient humidity and temperature. Latency time before feeding began, feeding activity, and weight change were recorded during the feeding trial. Latency time was determined by visually scanning all individuals each minute for the first 60 minutes of the feeding trial and recording when feeding began. After the first 60 minutes, each individual was assessed for feeding activity (yes or no) at 30-minute intervals. Each female was weighed before being introduced to the feeding trial and immediately after the feeding trial was concluded. Weight change was quantified as the percent of the initial body weight gained or lost per hour during the feeding trial. Trials began between 8:15 am and 12:06 pm PT and concluded between 3:12pm and 7:16 pm PT between January 28 and February 25, 2014, all during daylight or twilight hours when soapberry bugs normally feed.

After feeding trials concluded, females were returned to the growth chamber and retained with the seeds in their treatment pod (no additional seeds were added) for the remainder of the experiment. After a week, females were given an opportunity to mate. Mates were assigned from within the same population, but from different mothers, who had emerged at least 48 hours prior to introduction to allow for the complete hardening of the cuticle. If multiple suitable males were available, pairs were generated randomly. A single male was isolated with each female in a clean inverted plastic cup (45mm diameter, 31mm tall) within each female’s container for 24 hours to allow mating. The seedpod was excluded from the cup to stop males from feeding on treatment seeds. Once a female started laying eggs after mating, eggs were collected and weighed daily for the next 10 days, or until death. Individuals who did not lay eggs in the first 30 days after eclosion (the estimated adult life expectancy in the field (Carroll 1991)) were counted as not reproducing. At the end of 30 days, or after mortality, all individuals were frozen at −20°C and stored for morphological analyses. Closed seedpods were then opened and each seed was examined for feeding damage. Beak length, thorax width, and wing length were measured for all individuals with Mitutoyo calipers under a dissecting scope to ±0.02mm accuracy.

#### Field feeding trial

The third experiment (Fig. 2c) was designed to test the effect of the balloon vine seedpod on adult performance (H2) and natural selection on beak length (H3) in the field. The field feeding trial was conducted in Key Largo using adult females collected from the field during a 3-day period from April 6-8, 2014. Instead of dry pods, this experiment used mature green pods of *C. corindum* collected in the field containing naturally occurring seeds. In this experiment, the seedpod treatment was less invasive: for the ‘open’ treatment, all three segments of the seedpod were opened manually, and for the ‘closed’ treatment seedpods were left intact as they were found in the field. Individual females were introduced to mesh-lidded BugDorms with a single pod of their treatment group. Trials were conducted outdoors in Key Largo in the shade on April 9, 2014. Females were starved by withholding food (but not water) for 24 hours prior to the feeding trial to motivate feeding. Prior adult feeding and individual age were unknown. Individuals were introduced between 8:20 and 8:57 am and removed between 4:10 and 5:22 pm. Latency to feed and time spent feeding were measured for each individual. Beak length, thorax width, and wing length were measured alive for all individuals using Mitutoyo calipers.

## Statistical Analyses

All analyses were conducted in R version 3.2.2 (“Fire Safety”). The sets of models evaluated for each response variable are discussed in greater detail below. For each response variable, all models were compared using the Akaike Information Criterion (AIC), a metric that ranks the relative quality of a set of models based on fit and simplicity, with the exception of egg number, for which models were compared using weighted AIC (wAIC). All models with a >5% probability of being the best model out of the set were examined for each response variable. Effects are only reported for factors that were consistent in their effect direction and significance in all examined models unless otherwise stated. The specific test statistics and effect sizes reported in the results section were taken from the model with the highest probability.

### Statistical analyses of juvenile traits

For analyses of juvenile traits, the performance cost of the seed coat (H1) was evaluated by including the seed treatment as a fixed factor in the analyses of survival and development time. The developmental effect of each host plant on morphology (H2) was evaluated by including rearing host plant as a fixed factor in the analyses of beak length, body size, and wing morph. Recent work demonstrates that gene flow between host-associated populations is likely very high in the southern part of the range, particularly from the invasive to the native host (Cenzer 2016). Moving north away from where the two plants co-occur, rates of gene flow between the two hosts should decrease. Therefore, I evaluated the effect of gene flow on beak length plasticity (H3) by including the interaction between latitude and rearing host plant in the analyses of beak length. Analyses for each juvenile response variable are discussed in more detail below.

For each juvenile response variable (survival, development time, thorax width, beak length, and wing morph), all possible models including the main effects of rearing host (*C. corindum* or *K. elegans*), ancestral host (*C. corindum* or *K. elegans*), sex (male or female), and all possible two-way interactions were considered. All models were also analyzed as generalized linear mixed models with the additional random factor of individual population nested within collection host and the random factor of family nested within individual population. For survival, seed treatment (cracked or uncracked) was also included as a main effect and in all possible pair-wise interactions. Sex was excluded from the analyses of survival because it was not possible to sex individuals that died before reaching adulthood. For analyses of beak length, I used the residuals of the linear model log(beak length)~log(thorax width) as the response variable. This controlled for body size and improved both normality and homoscedasticity.

Differences in survival between treatments created heavily unequal sample sizes in all morphological traits and development time. Survival in high mortality treatments may have been non-random with respect to the genotypes that successfully reached adulthood, causing a single generation of strong selection. In order to distinguish plasticity from short-term selection, I only evaluated development time and morphological traits for individuals in the cracked seed treatment, which had high survival across host plants. Analyses of beak length revealed the unexpected result that plasticity in response to rearing host was maladaptive (Fig. 5a); this led to the hypothesis (H3) that maladaptive plasticity was the result of gene flow. To test H3, the top six models for beak length were re-analyzed with the added fixed factor of latitude and the latitude*rearing host interaction. I assessed whether or not the addition of latitude and the interaction improved model fit using Chi-squared tests comparing (1) each model with vs. without latitude and (2) each model with latitude vs. with the latitude*rearing host interaction. Latitude and the interaction were included post-hoc; therefore, the results of these comparisons are discussed here in the context of hypothesis development rather than hypothesis testing, and latitude was not added to the analyses of any other traits.

Survival and wing morph were modeled using a binomial error distribution, while development time, thorax width, and beak length had Gaussian error distributions. Five outliers, identified using the boxplot function in R, were excluded from the analyses of development time. The results were qualitatively the same if 3 of these outliers (45, 46, and 48 days) were included (mean±95% CI development time was 31.17±0.83 days); however, the analyses were not robust to the inclusion of two extreme outliers (72 and 73 days). These outliers were likely caused by individuals exhausting their food supply prior to reaching a large enough size to molt to adulthood, and are probably not informative for assessing any of the effects intended to be studied in this experiment. Normality of residuals was tested using Shapiro-Wilk tests and homoscedasticity was assessed using studentized Breusch-Pagan tests (in the R package lmtest).

### Statistical analyses of adult traits

For analyses of adult traits, the performance cost of the seedpod (H4) was evaluated by including the seedpod treatment as a fixed factor in the analyses of each response variable. Natural selection on beak length (H5) was evaluated in four ways: (1) the effect of beak length in lab trials on the response variables latency time and time spent feeding in the open pod treatment; (2) the effect of beak length in lab trials on the number of seeds fed on in closed pods; (3) the effect of the interaction between beak length and seedpod treatment on latency time and feeding activity in field trials; and (4) the effect of the interaction between beak length and seedpod treatment on egg production in lab trials. Analyses for each response variable are discussed in more detail below.

In lab feeding trials, 0/48 individuals were observed feeding through closed pods while 58/63 individuals were observed feeding on open pods; the effect of the seedpod treatment on latency to feed and feeding activity in the lab were considered to pass the intraocular trauma test and therefore did not undergo any formal analyses. I analyzed latency to feed and feeding activity in the lab on open pods only. For both of these response variables, I analyzed all possible generalized linear models including the main effects of host plant, beak length, and body size and all possible pairwise interactions. Family (nested within host plant) and individual (nested within family) were included as random factors in generalized linear mixed models. Latency time had a negative binomial error distribution and feeding activity (whether an individual was observed feeding at each observation time) had a binomial error distribution.

In field trials, I analyzed the effect of pod treatment, beak length, and body size on latency time and feeding activity. For both response variables, I tested all generalized linear models including pod treatment, beak length, body size, and all possible pairwise interactions. Field trials were conducted in Key Largo with naturally occurring pods; therefore only one host plant (*C. corindum*) was used. Generalized linear mixed models including individual number as a random factor were also analyzed for field trials. I analyzed field latency time with a Gaussian error distribution and field feeding activity with a binomial error distribution.

Weight change, egg production, egg weight, and the number of seeds consumed were only measured in the laboratory experiment. To explore what factors effected weight change, I analyzed all possible models including host, seedpod treatment, beak length, and all possible two-way interactions with a Gaussian error distribution. The effect of seedpod and host plant on individual egg weight was analyzed by comparing all possible models including treatment, host plant, and body size main effects and body size*host plant and body size*treatment interactions using a Gaussian error distribution. I treated whether or not each seed had been fed on as a binomial response variable and analyzed generalized linear models looking at the fixed effects of host plant, beak length, and the beak length*host plant interactions as well as the random effects of individual identity and family.

I analyzed egg production using Markov Chain Monte Carlo (MCMC) simulations in the ‘map2stan’ function in the ‘rethinking’ package in R (McElreath 2016). No individuals in the closed pod treatment on the host *C. corindum* produced any eggs. Generalized linear models cannot handle a treatment combination with no variance; MCMC was the best modeling approach for dealing with this problem. I approximated a negative binomial distribution for egg number using a gamma Poisson; the negative binomial distribution is commonly used for reproduction data because it is the distribution that results from many independent trials (in this case, females) where each trial is a series of yes/no events (in this case, whether or not an egg was produced at a given time point). Models were run for each main effect alone and for all additive and pairwise interactions. The model including all three main effects plus the treatment*beak length interaction was run to test for the effect of divergent natural selection on beak length by seedpod treatment with a host effect. All models were compared using weighted AIC using the ‘compare’ function in the rethinking package. One model held 100% of the weight; therefore results are reported only for this model.

## Results

### Assessing the effects of rearing host and seed coat on juveniles

#### Cracking the seed coat increases juvenile survival (H1)

The cracked seed treatment was associated with a dramatic increase in survival probability on both *C. corindum* (0.079 vs 0.94) and *K. elegans* (0.83 vs 0.98)(z-value=7.70, p<0.001) (Fig. 3). This was in addition to the clear difference in survival between the two rearing host plants (z-value=9.43, p<0.001) which was also observed in earlier experiments (Cenzer 2016). The effect of the cracked seed treatment increasing survival was stronger for individuals reared on *C. corindum* (z-value=−2.23, p=0.026). Nymph mortality was heavily skewed towards very young individuals, such that 94% of nymph mortality occurred in the first 7 days after hatching (Fig. S1).

**Figure 3.**
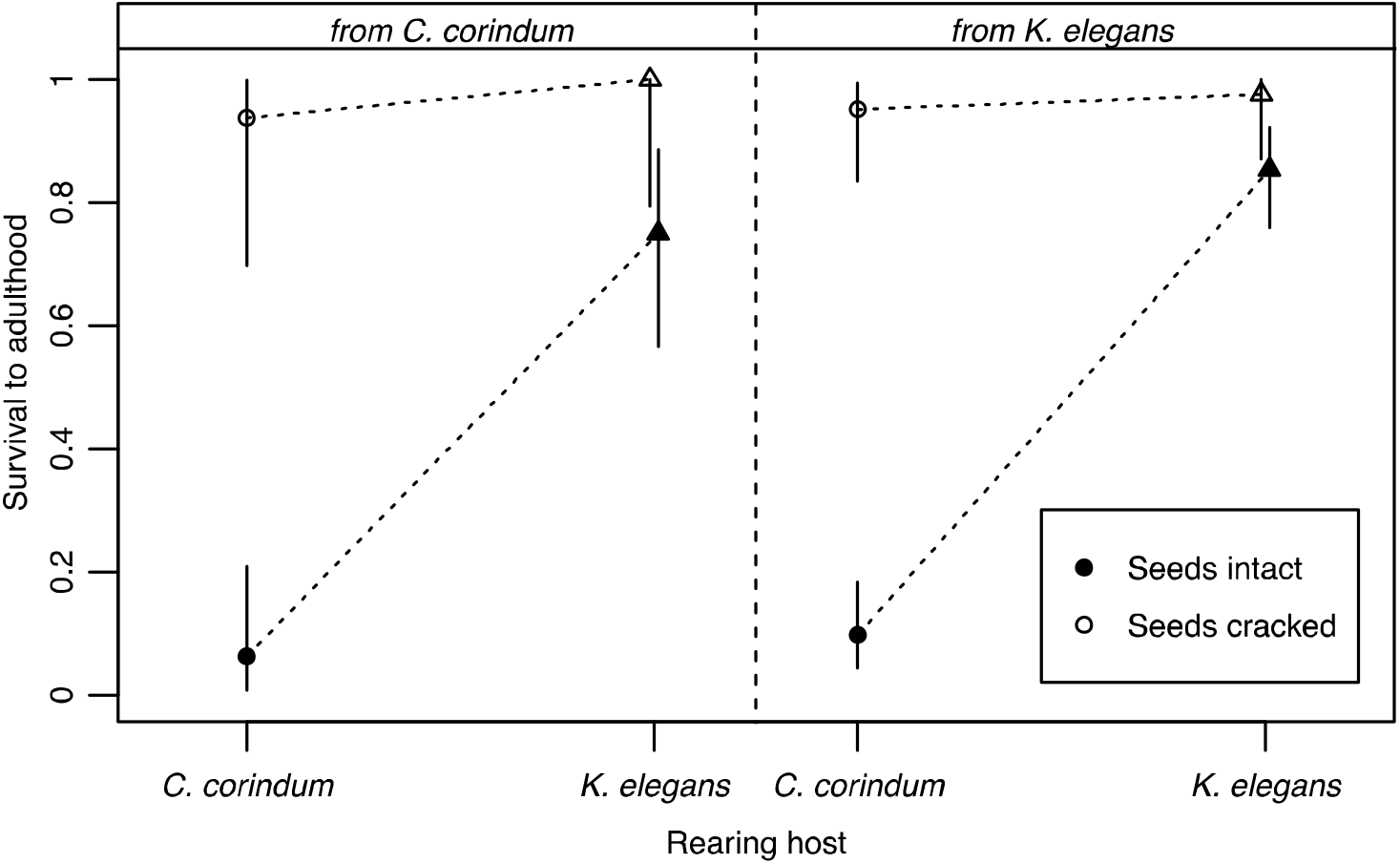
Soapberry bug juvenile survival on seeds with an intact (black) or a cracked (white) seed coats when raised on the native host, *C. corindum* (circles), or the invasive host, *K.elegans* (triangles). Points on the left represent bugs whose ancestors were collected on *C.corindum* while points to the right were collected from *K.elegans.* Error bars represent the 95% binomial confidence interval using the Pearson-Clopper method; points are jittered slightly for error bar visualization.

#### Rearing host has strong plastic effects on morphology & development time (H2 & H3)

There was a significant effect of rearing host on development time, such that individuals reared on *K. elegans* developed more slowly than individuals reared on *C. corindum* (t-value=3.35, p<0.001, df=98) (Fig. 4). Historically, development time was faster for populations from *K. elegans*; however, although the trend in this experiment was in that direction, it was not significant.

**Figure 4.**
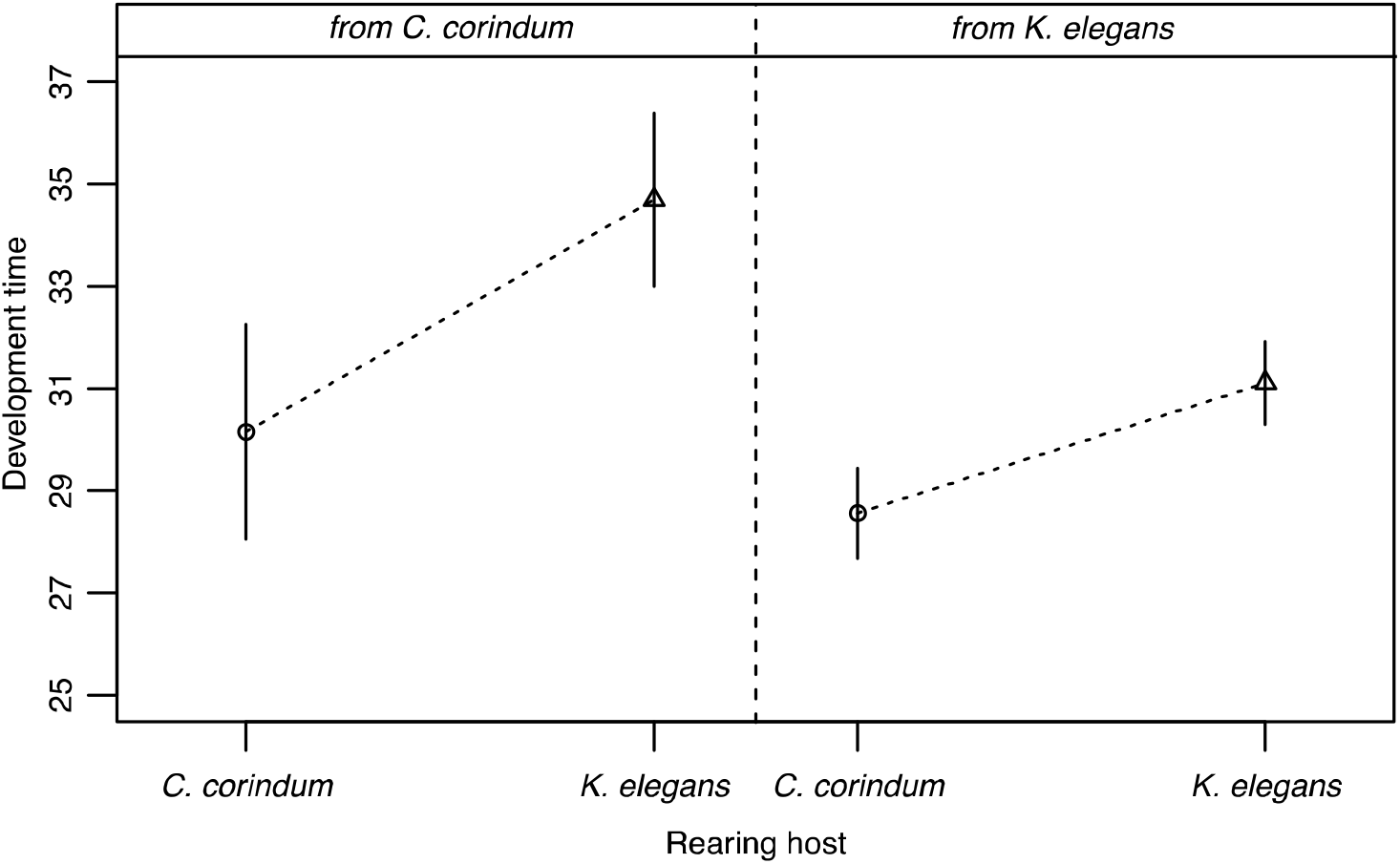
Soapberry bug juvenile development time on seeds with cracked seed coats when raised on the native host, *C. corindum* (circles), or the invasive host, *K. elegans* (triangles). Points on the left represent bugs whose ancestors were collected on *C. corindum* while points to the right were collected from *K. elegans.* Error bars represent 95% confidence intervals.

Individuals descended from populations on *K. elegans* had shorter beaks than those from *C. corindum* (t-value=−5.22, p<0.001, df=105)(Fig. 5a) as predicted by historical patterns of local adaptation (Carroll and Boyd 1992). The effect of rearing host on beak length was significant and in the opposite direction: individuals reared on *K. elegans* had longer beaks than those reared on *C. corindum* (t-value=2.79, p=0.006, df=105). Males had shorter beaks than females, consistent with known sexual dimorphism (t-value=−3.64, p<0.001, df=105). Adding the interaction between latitude and rearing host plant significantly improved the fit of all tested models. The effect of latitude alone was not significant (t=−0.36), but there was a significant interaction between latitude and rearing host plant. The effect of rearing host plant on beak length became weaker moving north away from where the two plants co-occur (t=−2.51, p=0.014, df=103).

**Figure 5.**
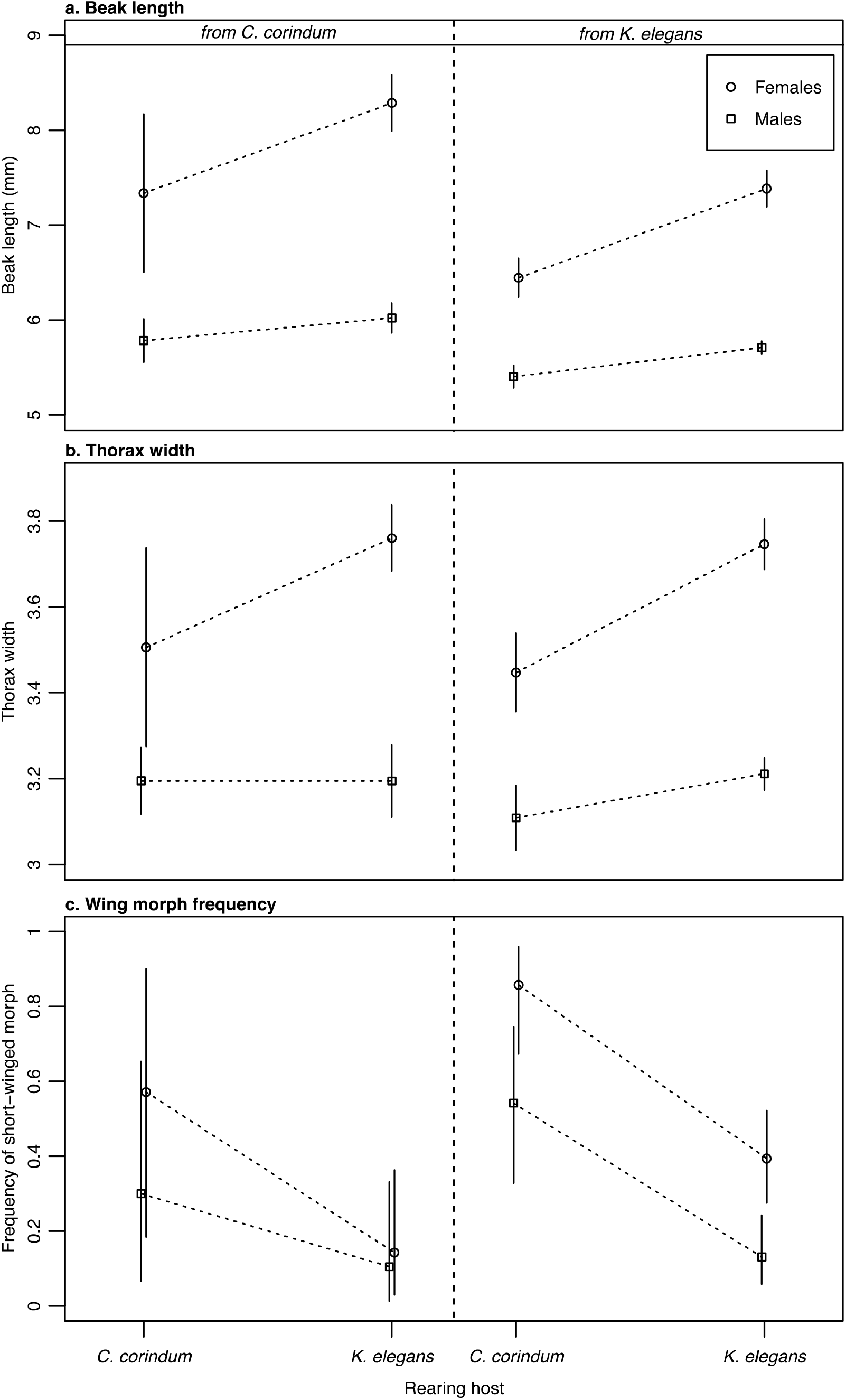
Adult morphology for females (circles) and males (squares) for bugs with ancestry from *C. corindum* (left panels) and *K. elegans* (right panels) when reared on *C. corindum* or *K. elegans* (x-axis, Rearing host). a. Beak length (mm). b. Thorax width (mm). c. Frequency of short-winged (brachypterous) wing morphs.

Individuals reared on *K. elegans* had significantly larger thorax widths than individuals reared on *C. corindum* (t-value=4.08, p<0.001, df=106) (Fig. 5b). Males had consistently smaller thorax widths than females (t-value=−5.93, p<0.001, df=106). There was a significant interaction such that males raised on *K. elegans* were smaller than predicted by the main effects alone (t-value=−2.21, p=0.03, df=106).

Being from a population collected from *K. elegans* increased the probability of having short forewings (z-value=−2.29, p=0.022). In contrast, being reared on *K. elegans* decreased the probability of having short forewings (z-value=4.51, p<0.001). Being male also decreased the probably of having short forewings (z-value=3.11, p=0.002)(Fig. 5c).

### Assessing the effects of the seedpod on adult performance

#### The host seedpod decreases adult performance (H4)

In field trials, 13 of the 15 individuals in the open treatment were observed feeding, while 4 of the 15 in the closed treatment were observed feeding (z=2.45, p=0.014). Individuals on the closed treatment took longer on average to begin feeding, although this effect was not statistically significant (Fig. 6b). Individuals in the closed pod treatment spent significantly less time feeding than individuals in the open treatment (z-value=3.63, p<0.001)(Fig. 6d). Results for beak length are discussed in the next section.

**Figure 6.**
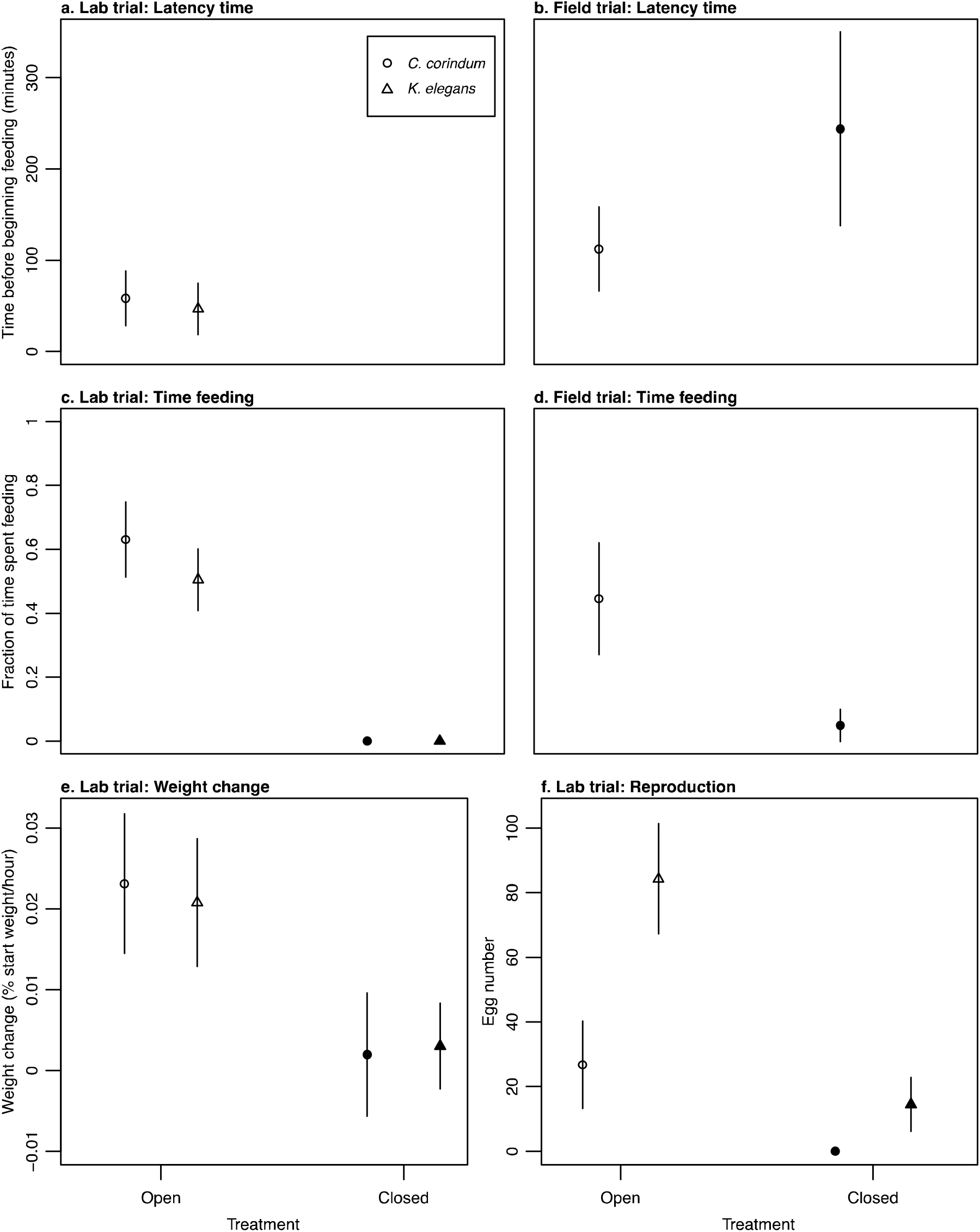
Performance metrics for open (white) and closed (black) pod treatments on *C. corindum* (circles) and *K. elegans* (triangles) in the lab (a, c, e & f) and field (b & d). a. Latency time before beginning feeding in lab trials. No individuals successfully fed through closed pods, so there is no latency time for those treatment groups. b. Latency time before beginning feeding in field trials; measures only collected on *C. corindum* due to the absence of naturally occurring closed pods on *K. elegans*. c. Fraction of time spent feeding in lab trials. d. Fraction of time spent feeding in the field trial on *C. corindum* only. e. Weight change as percent of initial body mass gained per hour in lab feeding trials. f. Number of eggs produced in first 10 days of reproduction in lab trials.

In lab trials, there was a clear and unambiguous effect of the seedpod on both latency time and time spent feeding (Fig. 6a & 6c). Individuals on *K. elegans* began feeding faster than individuals on *C. corindum* (z-value=−2.75, p=0.006)(Fig.6a) and individuals with larger bodies took longer to begin feeding overall (z-value=2.34, p=0.019). For feeding activity, individuals on *K. elegans* were observed feeding less frequently than those on *C. corindum* (z-value=−2.05, p=0.04)(Fig. 6c). Individuals in the open pod treatment gained more weight per hour than individuals in the closed pod treatment (t-value=4.757, p<0.001, df=92)(Fig. 6e). There was no significant effect of seedpod treatment on individual egg weight (t-value=−1.16, p=0.25). Larger females produced larger individual eggs (t-value=2.68, p=0.01, df=58). Females from *C. corindum* laid individual eggs 12% larger than those from *K. elegans* (t-value=−5.34, p<0.001, df=58) and had a stronger relationship between thorax width and egg size (t-value=2.87, p=0.006).

For egg production, the model including host species, seedpod treatment, and the interaction outperformed all other models (weight=1). Females laid more eggs in the open pod treatment than in the closed pod treatment (effect estimate 89% quantile: 6.22, 57.06)(Fig. 6f) and females on *K. elegans* laid more eggs than females on *C. corindum* (effect estimate 89% quantile: 5.59, 56.36). The interaction between pod treatment and host plant indicated that the positive effect of open pods on egg production was stronger on *C. corindum* (89% quantile: −54.80, −4.05); this is likely due to the fact that any increase from zero eggs would be proportionally larger than an increase from some eggs.

#### Beak length influences performance on open and closed seedpods (H5)

Data on latency time in both the lab and field and the number of seeds consumed in the lab support the hypothesis that the host seedpod contributes to divergent natural selection on beak length. In the field, individuals with shorter beaks began feeding more quickly than individuals with longer beaks in the open treatment (t-value=2.64, p=0.023, df=11)(Fig. S2). There was also a significant interaction between thorax width and treatment, such that larger bodied individuals in the open treatment began feeding more quickly than small-bodied individuals (t-value=−2.864, p=0.015, df=11). There was no significant effect of beak length or the beak length*treatment interaction on total time spent feeding in the field.

In the lab, individuals with longer beaks took longer to begin feeding when pods were open (z-value=1.98, p=0.048, df=44), consistent with results in the field. Unlike field results, larger individuals in the lab took longer to begin feeding (z-value=2.338, p=0.02). The combined effects of beak length and body size in the lab were less than additive; eg, the interaction between body size and beak length had a significant negative effect on latency time (z-value=−2.05, p=0.041). Individuals feeding on *K. elegans* showed a stronger effect of beak length on increasing latency time than those on *C. corindum* (z-value=2.56, p=0.011)(Fig. S2). Of the 48 individuals in the closed pod treatment in the lab, 12 individuals created damage on one or more seeds within the pod. There was a strong positive effect of beak length on the number of seeds damaged by an individual (z-value=2.32, p=0.02, df=118)(Fig. 7).

**Figure 7.**
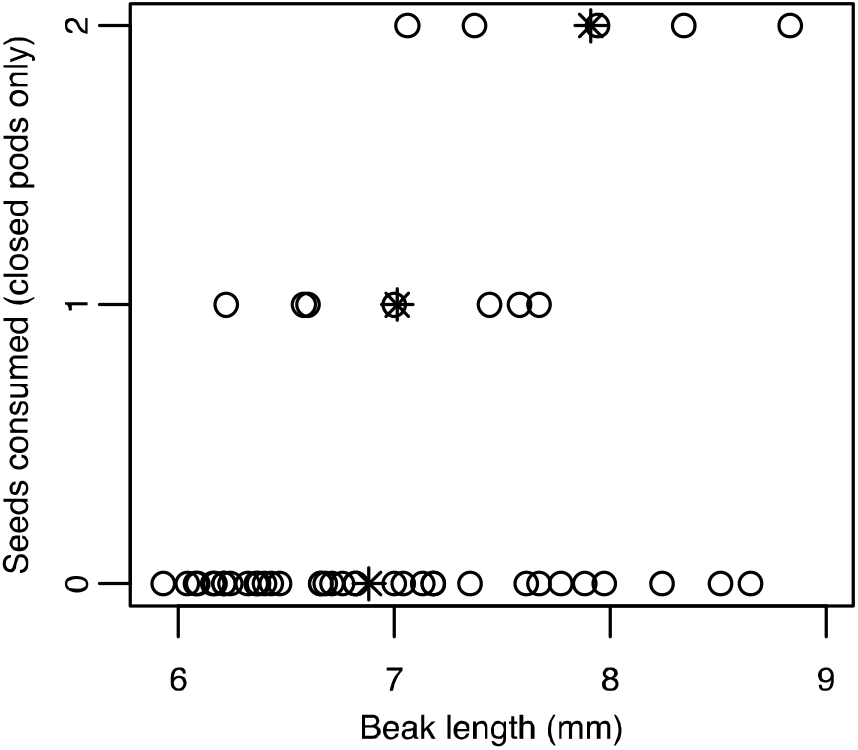
Number of seeds consumed in closed pods in lab feeding trials as predicted by individual beak length. Stars represent mean beak lengths that consumed 0, 1 or 2 seeds.

## Discussion

There are five main results of this work. First, the tough seed coat drives host-associated patterns of juvenile mortality in soapberry bugs. Second, the seedpod severely inhibits adult performance through feeding success, weight gain, and reproduction. Third, I found support for the hypothesis that the seedpod exerts divergent natural selection pressure on beak length: long beaks improve feeding success on the closed pods of the native *C. corindum* and short beaks improve feeding efficiency on the open pods of the introduced *K. elegans*. Fourth, developmental plasticity was maladaptive, causing females that were reared on each host plant to develop beaks mismatched to the pod size of their rearing host. Finally, plasticity was strongest where the two host plants occur in close proximity, consistent with maladaptive plasticity emerging as a result of ongoing gene flow between host plants. Taken together, these results indicate that the native host plant, *C. corindum*, has strong negative effects on juvenile fitness via the seed coat, strong negative effects on adult performance via the seedpod, and counteracts natural selection via maladaptive plasticity on adult morphology. This suggests that the native host has become a ‘fitness valley’ for soapberry bugs, resulting in much lower performance than the invasive *K. elegans*. Gene flow appears to be contributing to maladaptive plasticity in beak length that masks the genetic differences between host-associated populations in the field.

Juvenile survival, which was locally adapted between *K. elegans* and *C. corindum* in 1988, was dramatically reduced in subsequent decades, likely as a result of asymmetric gene flow from populations adapted to the invasive host to populations on the native host (Cenzer 2016). The proximate reason for juvenile mortality on the native host, and to a lesser extent on the introduced host, is demonstrated here to be the seed coat. Given the early mortality that occurs for individuals raised on intact seeds, this is probably the result of newly hatched individuals being unable to penetrate the seed and subsequently starving to death. It could also be that chemical defenses within the seed’s exterior, rather than or in addition to the physical barrier, are responsible for early mortality. This opens up further lines of inquiry in this system: Has the thickness or toughness of the seed coat on the native host changed during the last three decades? What aspects of nymphal morphology or physiology allow some individuals to deal with the tough seed coat while others do not? It is possible that this plant defense also exerts selection on beak length, but in first instars rather than in adults, which may be pleiotropically affected by the same factors that influence adult beak morphology.

The second plant defense that is clearly important for driving fitness in the field on the native host is the seedpod. The presence of a closed seedpod severely inhibited adult female performance: it negatively effected time to begin feeding, time spent feeding, weight gain, seed access, and egg production. The fitness costs of the seedpod make it a potentially potent driver of natural selection. Performance for short-beaked females in the field may not be as grim as measured in this study, however. Females that are incapable of feeding through closed seedpods in the field may instead compete with males, juveniles, and other short-beaked females for access to seeds after the pods dehisce. All life stages must also compete with other native seed predators on *C. corindum*, most of which feed prior to the opening of the seedpod (eg, the silver-banded hairstreak [*Chlorostrymon simaethis*], Carroll and Loye 2006). The seeds of the introduced host have not been adopted as a food source by any other species, so competition on this host is entirely intraspecific.

The pattern of beak length evolving to match local seedpod size has been observed in multiple populations and species of soapberry bugs across hosts and continents (Carroll and Loye 1987; Carroll and Boyd 1992; Carroll and Loye 2012). These patterns of local adaptation are highly suggestive of divergent selection being exerted by seedpod size on beak length. Carroll and Boyd (1992) suggested that individuals with longer beaks have a competitive advantage by being able to access seeds in large pods, while individuals with shorter beaks have an advantage of greater feeding efficiency on flattened pods. In this study, I found support for both of these hypotheses, although I did not find evidence that they scale up to measurable differences in reproduction. It may be that the advantages of increased feeding success and efficiency for fitness are relatively subtle or that they are only accrued in the presence of competition, which was not included in this study.

Rearing host plant had a strong plastic effect on beak length, body size, and wing morphology. On both hosts, females developed beak lengths that were mismatched with local pod size: females reared on *C. corindum* developed short beaks and females reared on *K. elegans* developed long beaks. This pattern of maladaptive plasticity was not observed for beak length in host races studied in the 1990s; however, it was observed in crosses *between* host races (Carroll et al. 2001). Since individuals with mixed ancestry expressed maladaptive plasticity in the 1990s, increasing gene flow could have increased maladaptive plasticity by creating hybrids and backcrosses in nature. More spatially separated populations should have lower rates of gene flow; therefore, we should expect that hybrids and backcrosses are more common in areas where the two hosts are close together. Consistent with this expectation, I found that developmental plasticity was strongest in the south where the ranges of the two host plants overlap and became weaker moving north away from the native host. The spatial pattern of beak length plasticity and the potential role of gene flow in generating that plasticity both warrant further study.

It is also possible that correlated selection on beak length has changed over the past several decades if selection pressure on wing morph or development time has changed. Developing on the introduced *K. elegans* caused individuals in this study to develop more slowly, achieve larger overall body sizes, and produce more dispersive morphs than developing on *C. corindum*. There is a genetic correlation between wing morphology and beak length (Dingle et al. 2009) and I documented a strong correlation between beak length and development time in this study (Fig. S4). If selection on wing morph plasticity or development time has changed since 1988, beak length plasticity could have evolved as a result of correlated selection. Understanding how selection is acting on dispersal morphology and development time in nature, and how that influences correlated selection on beak length, merits further investigation.

Maladaptive plasticity itself may exacerbate gene flow, as individuals dispersing from their natal host to the alternative host will have the advantage of a beak length that better matches local pod size. For example, soapberry bugs dispersing from *K. elegans* to *C. corindum* will have longer beaks because of their developmental environment and will therefore be more capable of accessing seeds in the large pods of *C. corindum* even though they are genetically ‘shorter-beaked’. Dispersing females from *K. elegans* will have the further advantage of increased reproductive output. However, their offspring will then be at a severe disadvantage when they develop on *C. corindum*, as their beaks will be shorter both genetically and plastically. The plastic effects of host plant on dispersal morphology should result in increased dispersal from *K. elegans* to *C. corindum* and reduced dispersal in the other direction, exacerbating gene flow that already appears to be asymmetric due to differences in host plant abundance.

Intuitively, maladaptive plasticity should be most common in scenarios where selection has not had an opportunity to act. Maladaptive plasticity has been observed in cases where organisms encounter novel environments that either expose cryptic variation or create a mismatch between perceived cues and the quality that cue is associated with in ancestral habitats (Schlaepfer et al. 2002; Ghalambor et al. 2007; Carroll 2008; Hale and Swearer 2016). Maladaptation may also arise without environmental change if the genetic combinations exposed to selection are themselves novel. This process is typified in hybrid zones, when haplotypes that have been diverging in isolation are combined following secondary contact to produce novel, usually detrimental, phenotypes in hybrids (Burke and Arnold 2001). In soapberry bugs, we see evidence of increased gene flow from populations on *K. elegans* to populations on *C. corindum* (Cenzer 2016). Although natural selection on beak length is ongoing in soapberry bugs, it appears to be unable to overcome the homogenizing effects of gene flow and plasticity. Although genetic differences in beak length remain between host-associated populations, they are entirely masked in the field by the opposing effect of maladaptive plasticity.

## Acknowledgements

I would like to thank R. Brennan, P. Brand, J. Berg, B. Cornwell, S. Ehlman, L. Yang, A. Sih, S. Carroll, and J. Rosenheim for comments on earlier drafts of this manuscript. I would like to thank A. Sharma for assistance with the juvenile rearing experiment. I would also like to thank the Florida Parks Department for permission to collect *J. haematoloma* and *C. corindum* seeds within Florida State Parks.

